# AI-Enhanced Marker-Assisted Selection Concept for The Multifunctional Honey Bee (Hymenoptera: Apidea) Protein Vitellogenin (Vg)

**DOI:** 10.1101/2025.05.13.653757

**Authors:** Vilde Leipart, Gro V. Amdam, Sharon O’Brien, Elisabeth Pigott, Garrett Dodds, Kate E. Ihle

## Abstract

Managed honey bees (Hymenoptera: Apidae: *Apis mellifera* L) have experienced unsustainably high rates of annual loss driven by several interacting factors, most notably pests, pathogens, pesticides, and poor nutrition. Breeding bee stocks that can cope with these challenges is a priority. Advanced molecular methods (marker-assisted selection, MAS) have enhanced the breeding efficiency of domesticated animals in recent years, but have not contributed strongly to honey bee stock improvements. This is largely because desirable traits of bees usually emerge from collective phenotypes of workers (sterile females) instead of from the breeding individuals (queens and male drones). For collective phenotypes, single genes typically have small, additive effects, so identifying impactful MAS targets is challenging. Here, we provide proof of concept for a new approach to honey bee breeding through MAS using the multifunctional protein Vitellogenin (Vg), a protein known to interact with and mitigate the primary drivers of colony loss. Our pipeline leverages cutting-edge, artificial intelligence (AI)-driven protein structure modeling algorithms to predict the effects of genetic variants of Vg on relevant molecular functions including lipid, zinc, and DNA binding. Following the AI-powered Vg variant selection step, we use a combination of standard apicultural techniques and DNA sequencing validation to breed honey bee queens homozygous for the desirable Vg allele. Our protocol can kick-start a new area of modernized bee breeding: an AI-enhanced MAS system that allows cost-effective and nimble development of stocks to meet urgent and long-term needs of stakeholders.

## Introduction

Managed honey bee (Hymenoptera: Apidae: *Apis mellifera* L) colonies are an essential part of our agricultural landscape, and the pollination services they provide have an annual estimated economic value of nearly $15 billion in the United States alone (Bruckner et al., 2023). Domestic and international food supply depends on the timely availability of honey bee colonies for optimal production, but beekeepers are averaging colony losses at rates twice as high as historical records (Aurell et al., 2024). This situation threatens the beekeeping industry as well as global food security. Much research is, therefore, aimed at understanding and potentially reducing colony losses. However, we are learning from these studies that the insults to managed honey bee populations are many and interacting, and simple solutions are unlikely to materialize (French et al., 2024; Hsieh and Dolezal, 2024). Four generally agreed-upon factors drive the losses of managed honey bees: pests, pathogens, pesticides, and poor nutrition (DeGrandi-Hoffman and Chen, 2015; Dolezal et al., 2019; Dolezal and Toth, 2018). Their impacts are expected to worsen with the increased global temperatures and more unstable weather conditions that characterize climate change, as well as with increased globalization that involves more frequent introduction of pests and pathogens and more expansive use of new pesticides (Cornelissen et al., 2019; Insolia et al., 2022; Switanek et al., 2017; Zapata-Hernández et al., 2024). In this context, the future outlook for essential agricultural pollination services is increasingly grim. Marker-assisted selection (MAS) breeding can be a sustainable method to combat the effects of disease and other stressors. It is routinely used to adapt domesticated animals to changing expectations, such as modernized production or performance breeding standards, disease resilience, and improved feed utilization (Hammond, 1947; Olesen et al., 2000). In most production animals, breeding has progressed beyond first-generation breeding technologies based on individual trait phenotyping to second and third-generation technologies using MAS to identify breeding individuals more efficiently.

The domesticated honey bee is an excellent candidate for MAS. This is because of an unusually high recombination rate in the bee that breaks up associations between uninformative DNA markers and individual traits, leading to potentially rapid identification of genomic regions with strong influences on traits of interest (Beye et al., 2006). Yet, the selection of managed honey bees has not effectively progressed beyond basic colony- and queen-level traits, which are not sufficiently responsive to the challenges that face commercial pollination in the food supply chain. One explanation for this unfortunate situation is that some of the most valuable traits of honey bees are social traits, and it has proven difficult to find strong molecular markers that can assist the selection of social phenotypes (Behrens et al., 2011; Lapidge et al., 2002; Tsuruda et al., 2012).

Marker identification in animal breeding is typically a forward (molecular) approach, where researchers mine associations between available trait variation and massive data matrixes from technologies like full genome sequencing, transcriptomics, or proteomics. Alternatively, on can take a reverse approach to focus on promising candidate genes. In this context, we recently discovered over 100 variants of a single, health-promoting protein in honey bees, called Vitellogenin (Vg) (Leipart et al., 2022a). Vg is a 700 million-year-old protein fundamental to reproductive physiology in almost all egg-laying species. More comprehensively, Vg is a very high-density glycolipophosphoprotein with critical roles in lipid transport and Zn-binding. In honey bees, Vg takes on colony-level functions as a nutrient storage protein in workers and as a source of amino acids and other factors, including immune elicitors in the royal jelly that is synthesized by specialized nurse bee workers for transfer to colony members, including larvae and queen (Harwood et al., 2019; Seehuus et al., 2007). When Vg levels drop in nurses, this regulates the behavioral transition from inside nursing tasks to outside foraging tasks (Ihle et al., 2010; Marco Antonio et al., 2008; Nelson et al., 2007) and affects the preferences of foragers for collection of pollen or nectar (Ihle et al., 2010; Nelson et al., 2007). Vg additionally influences honey bee immunity in several ways, including as a transporter for the transgenerational immune priming of larvae, as a support for immune cell survival, and with roles in parasitic *Varroa* mite reproduction (Amdam et al., 2004; Dahlgren et al., 2012; Döke et al., 2015). More results also point to effects on resilience to wintering, oxidative stress and insecticides (Amdam et al., 2004; Döke et al., 2015; Fent et al., 2020; Nelson et al., 2007; Seehuus et al., 2006). Overall, it is thus not surprising that Vg is recognized by the scientific community and by stakeholders as a molecule tightly linked to honey bee colony health (Aurori et al., 2014; Smart et al., 2016).

The multiple roles of Vg were identified by reverse genetics, and our work has been central to advancing this knowledge: We successfully pioneered Vg gene knockdown technology for honey bees (Amdam et al., 2003; Guidugli et al., 2004; Nelson et al., 2007), representing the first successful gene knockdown in adult bee workers. Since then, our in-depth functional studies of Vg (Harwood et al., 2019; Ihle et al., 2010; Nilsen et al., 2011; Nunes et al., 2013; Seehuus et al., 2006), as well as follow-on work by others, (Christen et al., 2019; Fent et al., 2020; Kim et al., 2023; Ramsey et al., 2019) have revealed and cemented that Vg is a multi-functional protein – a Swiss Army knife – with a long range of structure-function relationships that can be targeted for selection. In fact, due to its diverse functions, Vg interacts with and may mitigate all the major factors that drive the recent colony losses with implications for honey bee nutrient storage, stress resistance and transgenerational immunity, as well as parasitic *Varroa* mite reproduction (Amdam et al., 2005, 2004; Harwood et al., 2019; Ramsey et al., 2019; Seehuus et al., 2006).

Recently, we predicted the structural and functional impact of genetic variation in Vg using the AlphaFold 2 algorithm (AF2, 2021). AF2 marked a sea change in protein structure biology by using deep learning based artificial intelligence (AI) to enable accurate protein structure predictions from protein sequences comparable to experimental methods, such as X-ray crystallography and nuclear magnetic resonance spectroscopy (Jumper et al., 2021). We used this technology to develop the first complete model of Vg, which was verified through electron microscopy of purified honey bee Vg protein (Leipart et al., 2022b). This complete model, combined with our knowledge about the specific Vg alleles of several commercially available honey bee stocks (Leipart et al., 2022a), sets the stage for a novel approach: Using the AlphaFold structural model to identify the most promising allelic selection targets to fast-track MAS for honey bees (Fig. 1).

**Figure 1:**
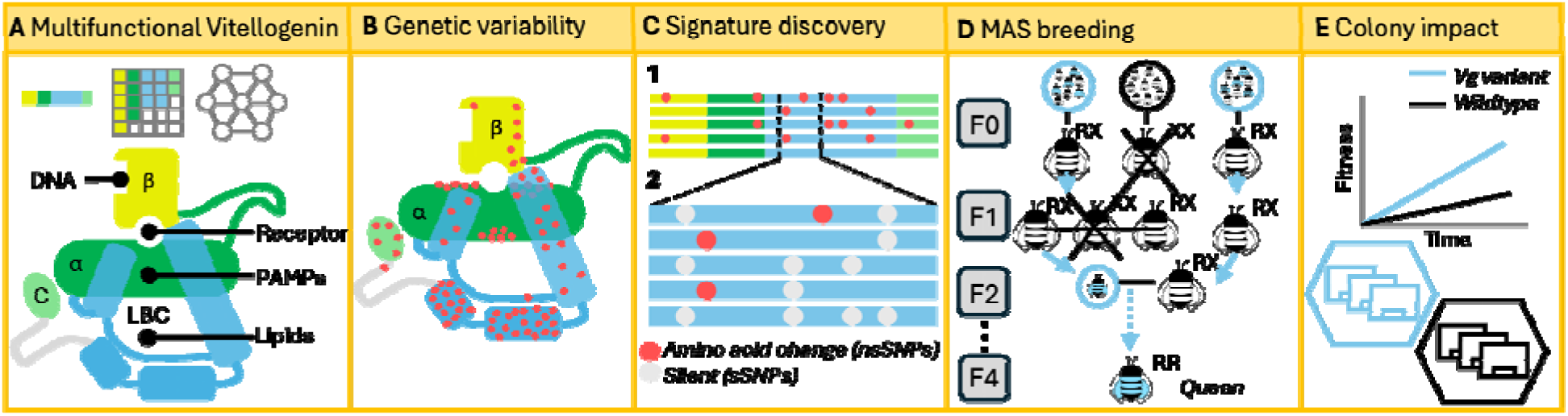
A) Honey bee Vitellogenin (Vg), predicted using AlphaFold consists of the β-barrel domain (β, yellow), the [7l-helical domain ([7l, green), the lipid-binding cavity (LBC, blue) and the C-terminal domain (C, light green). The functional sites for binding DNA, the receptor, pathogen-associated molecular patterns (PAMPs) and lipids are labelled. B) In the AlphaFold structural model, the identified nsSNPs (red dots) are located in the domains associated with the functional sites, which makes the potential targets for Vg MAS extensive. C.1) Long-range DNA sequencing determines Vg variants’ current diversity and availability in the selected colonies and identifies queens possessing at least one copy of the targeted Vg allele. C.2) The information from the long-range DNA sequencing is used to discover a “molecular signature” of Vg alleles that result in the protein variants of interest. D) The signature enables us to use standard, inexpensive Sanger sequencing to screen queens and drones to breed a population of bees homozygous for the targeted Vg allele (blue circles for drones, blue bees for queens) after a few generations (F4). E) Our approach can rapidly produce honey bee stocks with a desirable Vg variant to boost the colony’s fitness.

Thus, we here present a proof-of-concept that leverages the power of AI-enhanced structural predictions to facilitate MAS breeding of honey bee stocks. Our first-of-its-kind breeding scheme (Fig. 1) builds on high-throughput DNA sequencing and high-powered AI-assisted structural prediction model to identify Vg variants with predicted differences in structure-function relationships. Here, we target the Pol-line genetic stock, as our recent study revealed that this population has several variants with predicted functional differences, as well as a “reference” allele identical to the Vg variant present in the published honey bee genome (Leipart et al., 2022a). In addition, the Pol-line is well suited for a controlled MAS experiment since 1) this stock is a closed breeding population, which limits background genetic variation, and 2) the stock was selected to be highly Varroa resistant, improving survivorship. This last trait is especially important due to the more fragile nature of the single drone inseminated (SDI) colonies used to establish the Vg variant lines in our breeding program (see Materials and Methods).

In this proof of concept, we targeted reference Vg to demonstrate the combination of cutting edge AI-enabled proteomics, standard molecular tools, and instrumental insemination of queens can rapidly produce honey bee stocks with desirable protein variants (as outlined in Fig 1). This pipeline represents a new approach to solving the challenge of breeding honey bees to provide sustainable pollination services for the future.

## Material and Methods

### Initial screening bees

We previously identified 121 unique Vg molecules, (Vg protein variants) that differed in one or more amino acids (12) using Oxford Nanopore technology, which allows for labeling each amplicon with barcoded primers. By projecting the 121 Vg variants we identified onto a single molecular model of honey bee Vg, we found that they emerge from combinations of 81nsSNPs (12). The barcoding, moreover, enabled us to track each variant back to the country, apiary, colony, and individual bee.

Using this database, we identified domestic honey bee populations in the United States with Vg variants identical to the Vg reference published in the Honey Bee Genome initiative (NCBI Gene ID: 406088). The Minnesota Hygienic and Pol-line stocks, both bred to be resistant to Varroa mites, had relatively high occurrences of the reference variants 104, 107, and 111 in the initial broad sequencing effort (12). In the summer of 2023, we performed a more in-depth screening of these populations along with a third mite-resistant stock, Russian honey bees, not included in our original database. In total, we sequenced the Vg alleles of 416 haploid drones from 12 Minnesota Hygienic, 25 Pol-line, and 15 Russian honey bee colonies. Minnesota Hygienic bees were obtained from a single apiary in West Monroe, LA, USA. Russian and Pol-line samples were obtained from USDA-ARS apiaries in Baton Rouge, LA, USA.

Nearly all Pol-line colonies sequenced in 2023 were lost due to environmental exposure to toxic chemicals in Baton Rouge, LA USA. In the spring of 2024 we sampled a further 42 Pol-line colonies headed by queens that were either the daughters or sisters of queens from the original Pol-line sample.

### Nanopore DNA sequencing

Nine late-stage drone pupae were collected from the targeted colonies (52) in spring of 2023 onto dry ice and stored at −80°C until processing. Honey bee drones are haploid and have only a single set of chromosomes from their mother queen. Thus, fewer samples are needed to determine the genotype of the queen. We removed the head, abdomen and extremities from the thorax, placed the thoraxes into 2mL screw-cap cryo-vials, and filled the vials with 95% ethanol.

The samples were shipped to the Centre for Integrative Genetics (CIGENE) at the Norwegian University of Life Sciences in Norway for gDNA extracting, PCR, amplicon Oxford Nanopore Sequencing, and bioinformatic processing. The protocol used is described previously (Leipart et al., 2022a), but we applied a stricter threshold for the consensus sequences since this round only sequenced haploid drones (male honey bees): We only allowed for consensus sequences that were generated from single-haplotype samples (where the bioinformatic processing identifies only one type of raw read per sample). Applying the strict threshold and the error rate, as before, reduced our dataset to 330 samples. The mean number of raw reads per consensus sequence was 11602.72 (± 9611.947).

The consensus sequences were aligned to the reference sequence (NCBI Gene ID: 406088) using MAFFT (Katoh et al., 2019) to identify the start and stop codons in the Vg gene. The non-coding regions (before the start codon, after the stop codon, and intronic regions) were removed using JalView (v. 2.11.3.3) (Waterhouse et al., 2009). The coding region was translated to protein sequence and nsSNPs identified using MEGA-X (v. 10.2.4) (Kumar et al., 2018). We identified 26 different Vg variants (Table S1) that resulted from unique combinations of 34 nsSNPs.

### Signature discovery

We identified a Vg variant identical to the reference Vg (NCBI Gene ID: 406088) in 24 drones (Vg Variant 1, Table S1), 20 of which were from the Pol-line breeding stock. The 20 sequences were aligned to all sequences from the Pol breeding stock (141 sequences), using Jalview, to identify sites that uniquely separated the reference Vg from the rest. We identified 7 unique haplotypes (sequences with a unique combination of synonymous and non-synonymous SNPs) of exon 2 among the Pol-line drones (Fig S1). Haplotype 1 is identical to reference Vg, while haplotype 2-7 has an alternative nucleotide on one or several of the 9 SNPs sites along the exon. Together, the 9 positions create our signature for reference Vg, where we can differentiate the reference Vg haplotype from the other haplotypes in the Pol-line. The shorter sequence of exon 2 allowed us to use the more affordable Sanger sequencing to identify the reference Vg.

We also identified a second signature to target a site in the lipid-binding cavity of Vg. The Vg variant 15 (Table S1) includes the nsSNP p.S1110T. The nsSNP is positioned in the lipid-binding cavity with close contact (<6Å) with 3 phospholipids and 1 hydrocarbon chain. The lipid molecules from the crystal structure of lamprey Vg (Anderson et al., 1998) were inserted into the AlphaFold honey bee Vg model using TM-align (Zhang and Skolnick, 2005) and the distance was measured using PyMol (Schrödinger, n.d.). The nsSNP is predicted to contract the lipid-binding cavity by 146.016 Å^3^, using the variant predictor Missense3D (Ittisoponpisan et al., 2019). Taken together, the close contact with lipid molecules and predicted alteration of the cavity makes p.S1110T an potential nutritional-altering Vg target. Vg variant 15 includes 7 other nsSNPs, but they are not expected to have observable impacts on the lipid-binding properties of the cavity (Leipart et al., 2022a, p. 121). We used the same procedure, as described for the reference, to discover a short signature for Vg variant 15. Of the 92 drones with Vg variant 15, 48 were from the Pol-line breeding stock. Aligning the 48 alleles to all the sequences from Pol-line breeding, we identified a signature for Vg variant 15 in exon 4. In this exon region, we found 3 haplotypes resulting from 6 SNPs (Fig S2). Haplotype 1 is identical to reference Vg, while haplotype 2 and 3 has alternative nucleotides at several of the 6 SNPs in the exon 4 region. Haplotype 3 was found in 50 pol-line drones, 48 of them had Vg variant 15, while 2 had Vg variant 7. Vg variant 7 is rare and was only found for those 2 drones, collected from a single colony (P7). Vg variant 15 was prevalent in the Pol-line breeding stock and was identified in 16 colonies. Using the short sequence in exon 4 and excluding P7 from the selection, we can differentiate Vg variant 15 from other haplotypes in the Pol-line.

### Sanger sequencing

Primers for the reference signature were designed to cover nearly all of Vg exon 2 using Primer 3 (Kõressaar et al., 2018) and Primer-blast (Ye et al., 2012). A large fragment of Vg exon 2 was amplified with forward primer 5’ – GGGACAGTTTCAGCCGACTT-3’ and reverse primer 5’-TCTTGATCACCTCCATGTGGC-3’. A 620 base-pair length fragment was amplified via PCR with the program 2 min at 95° C followed by 35 cycles of 30 s at 95°C, 30s at 55°C, and 1min at 72°C. We confirmed fragment size and specificity on a 2% agarose gel. DNA for Sanger sequencing was extracted from wing clips from the queens and from thorax tissue from the drones. A similar procedure was followed for the lipid binding variant 15. We amplified a 542 bp fragment from Vg exon 4 with primers F: 5’ - AGTTTGATGAAGCTGAAGAGCC-3’ and R: 5’ -TTCCTTCCAGAGGAACGAGC-3’ and a 400 bp fragment from Vg exon 5 using primers F: 5’ -TGGACCAGAAGCCGAAGATG-3” and R: 5’ -TTAATCCTCGTAGAATACGTTGTTA-3’.

Wing clips were stored at −20° C until DNA was extracted. DNA extractions from queen wing clips were conducted using Zymo Quick-DNA Tissue/Insect Microprep Kit (N = 204). Queen wing samples were homogenized with 750 μL of Bashing Bead Buffer (Zymo Research, Irvine, CA, USA) using the OMNI Bead Rupter Elite Bead Mill Homogenizer (OMNI International, Kennesaw, GA, USA). Samples were homogenized four times at 4 m/s for 5 seconds. The DNA was extracted per the manufacturer’s protocol, including the optional step of adding β-mercaptoethanol to the Genomic Lysis Buffer. PCR was performed on the extracted DNA for Vg Exon 2. PCR products were purified using Qiagen QIAquick 96 PCR Purification Kit (Qiagen, Hilden Germany), following the manufacturer’s protocol. An aliquot of 10 µL per sample was then pipetted into a 96-well plate, sealed, and shipped at −20° C to the University of Illinois of Illinois Roy J. Carter Biotechnology Center Core Facility for Sanger sequencing reaction, column purification and electrophoresis on the AB 3730xl platform.

Drones were collected in to 2 mL bead tubes and stored at −20° C until DNA was extracted. Drones were homogenized using 600 μL of CTAB buffer using the OMNI Bead Rupter Elite Bead Mill Homogenizer for two cycles at 5 m/s for 12 seconds. DNA extractions were conducted using Promega Maxwell RSC PureFood GMO and Authentication Kit (Promega, Madison, WI, USA) per the manufacturer’s protocol for meat sample lysis. PCR was performed on the extracted DNA for Vg Exon 2. PCR products were purified using Qiagen QIAquick 96 PCR Purification Kit, following the manufacturer’s protocol. An aliquot of 10 µL per sample was then pipetted into a 96-well plate, sealed, and shipped at −20° C to the University of Illinois Roy J. Carter Biotechnology Center Core DNA Sequencing Facility for Sanger Sequencing. We genotyped 347 virgin queens and 497 drones using this DNA Sanger sequencing protocol.

### Queen rearing and insemination

We inseminated queens with at least one copy of the Vg reference variant with semen from a single drone from a mother queen hetero-or homozygous for the Vg reference variant. Those queens confirmed to be inseminated by a drone with the reference variant were used as breeder queens in the subsequent generations. This allowed us to significantly reduce generation time to just over a single month.

To produce new queens from mothers identified as having the reference signature, we grafted first instar larvae into plastic JzBz “queen cups” containing a sterilized royal jelly dilution, and introduced them to cell builders, colonies with a large population of young bees isolated from a queen, according to standard practice. Drawn queen cells were placed into glass vials in an incubator kept at 34°C. Each day, emerging queens were collected and individually tagged. approximately 2/3 of one wing were clipped for DNA analysis, and queens were introduced to a queen “bank,” a queen-less colony that sustains many queens kept in a single frame containing many small individual cages approximately 1” (2.54cm) in diameter and ¾” (1.9cm) deep for 7-10 days before insemination with semen from a single drone: approximately 1-2 microliters of semen depending on the drone, and 2 microliters of diluent. Pipette tips were changed and cleaned with a bleach solution in between inseminations to prevent cross-contamination.

Instrumentally inseminated queens were exposed to CO_2_ for approximately 3-5 min during the insemination and “banked” until DNA results were available. Then selected queens were given a second dose of CO_2_ and introduced into queen-less colonies; first in a small JzBz queen cage for five days, followed by transfer to a Scalvini cage until the queen began to lay eggs. Queens were then released freely into the colony. Grafting for the next generation of queens took place at least 7 days after the queen was confirmed to be laying fertilized eggs in the colony. We inseminated 98 of the screened queens.

## Results

### Initial screening for honey bee colonies with Vg reference variant

To evaluate the potential effect of Vg molecules, we need a reference (control) stock with a standard genetic background for future comparisons or relative scoring of the variant stocks. Therefore, we first aimed to establish a reference Vg stock of honey bees. Ideally, this stock can be used as a control to measure the effect of single amino acid changes on Vg, which can be used when evaluating production—and health-related trait differences between honey bee stocks.

The Vg variants published by Leipart et al. 2022 (Leipart et al., 2022a) were labeled during sequencing, allowing us to map the honey bee workers back to their original colonies. In this dataset, we identified three colonies with high incidences of reference variants in the worker bees (59-73%, Fig 2A). We selected 400 honey bee males (drones) from the three genetic stocks to screen for the reference Vg allele. The long-range DNA sequencing screen identified 20 drones from the Pol-line breeding stock that is identical to the reference sequence of honey bee Vg (NCBI Gene ID: 406088) (Fig 2B, Table S1). In addition to the reference stock, we also identified 48 drones with a lipid-binding cavity-altering amino acid change.

**Figure 2:**
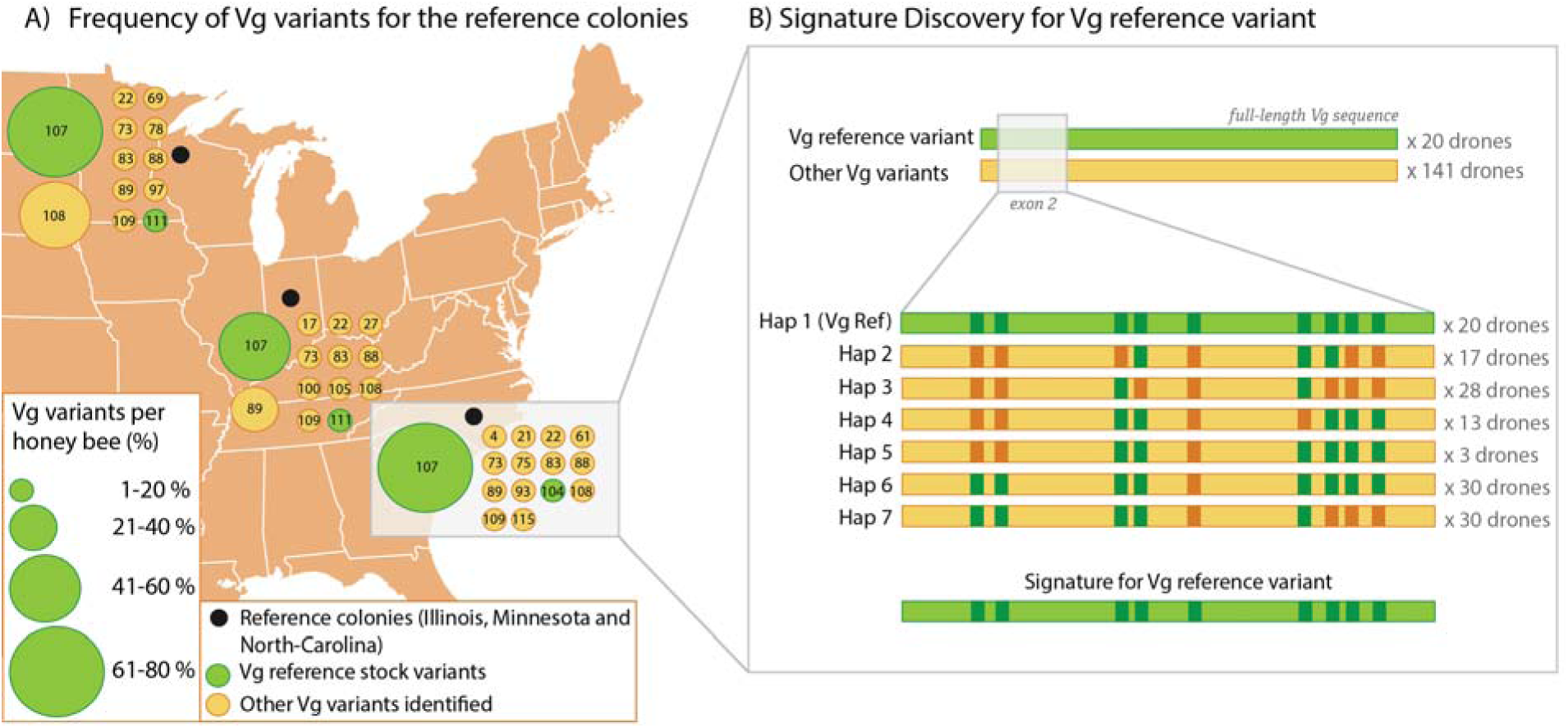
A) The Vg variants (numbered dots) identified at the reference colonies (black dots). The size and colors of the Vg variants illustrate the frequency and predicted impact of the identified nsSNPs, respectively. The genetic stock used in this study is marked with a grey box. B) In the selected stock, we identified 20 reference variants of Vg (green), while 141 other Vg variants were identified (yellow). The grey box marks the exon 2 sequence, zoom in below. Here we identified 9 SNPs that separate the reference Vg from the other Vg variants. SNPs with reference alleles are shown in dark green, while dark yellow SNPs have an alternative allele for each of the 7 unique combinations (haplotypes, Hap). The number of drones per haplotype is shown to the right. The signature for the Vg reference is shown on the bottom. For more details on specific SNPs sites and alternative nucleotides see Figure S1.

### Signature Discovery

We established a breeding scheme, explained in the next section, to create homozygotic queens for any Vg variant. To verify successful breeding, meaning that the offspring inherited the targeted Vg allele, we identified a “signature” for the targeted allele. The initial round of long-range Nanopore DNA sequencing allowed us to identify the Vg diversity in the colony. We used this information to establish the signature for the targeted Vg allele: a shorter sequence of the Vg gene where the targeted Vg allele has the same nucleotide at several polymorphic sites compared to every other sequence in that colony (Fig 2B). The unique combination of the SNPs was used to determine a Vg signature. We used our dataset to identify two signatures, one for the reference Vg (Vg variant 1) and one for a Vg variant with an amino acid change predicted to alter the lipid-binding cavity (Vg variant 15). Specifically for the reference Vg, we identified 9 SNPs in Vg exon 2, creating a unique signature (Fig 2B, see Figure S1 for nucleotide and position information) for the 20 drones at the Pol-line breeding stock.

We identified 6 SNPs in exon 4 for the lipid-altering Vg variant, creating a signature for the 48 drones (Figure S2). Two additional drones had the same signature in exon 4, which was genotyped to Vg variant 7, meaning that the drones have additional SNPs at positions outside of exon 4. However, Vg variant 7 was only identified in these two drones and found in a single colony (P7). Therefore, we could use the region in exon 4 to identify Vg variant 15 by avoiding sampling from colony P7.

In the following steps, we created a targeted Vg allele breeding stock. The short signature sequence (719 bp for the reference Vg and 416 bp for the lipid-altering Vg variant, compared to the full-length Vg of 6109 bp) allowed us to use a more efficient sequencing protocol (Sanger sequencing) to confirm the inheritance of the targeted Vg allele in queens and drones. To prove our concept, we created colonies with homozygotic queens for the Vg reference allele.

### Breeding scheme

In the summer of 2024, the reference signature allowed us to rapidly screen the surviving selected colonies, as well as daughter and sister colonies of the originally selected queens via Sanger sequencing of the 719bp Vg exon 2. We then again produced daughter queens from those queens found to have the reference allele. Wing clips from these queens were sequenced, and those with the reference allele were inseminated with semen from a single drone from a mother queen known to have a reference Vg allele. The thoraxes from these drones were then further screened for the reference signature via Sanger sequencing.

Those queens that were inseminated with semen from drones not possessing a reference allele were culled from the population. This process was repeated a second time to produce 35 colonies with workers homozygous for Vg reference alleles (Fig 3). The next generation will allow us to pool semen from multiple drones for more robust colonies. We are currently in the process of repeating this procedure for the lipid binding allele variants.

**Figure 3:**
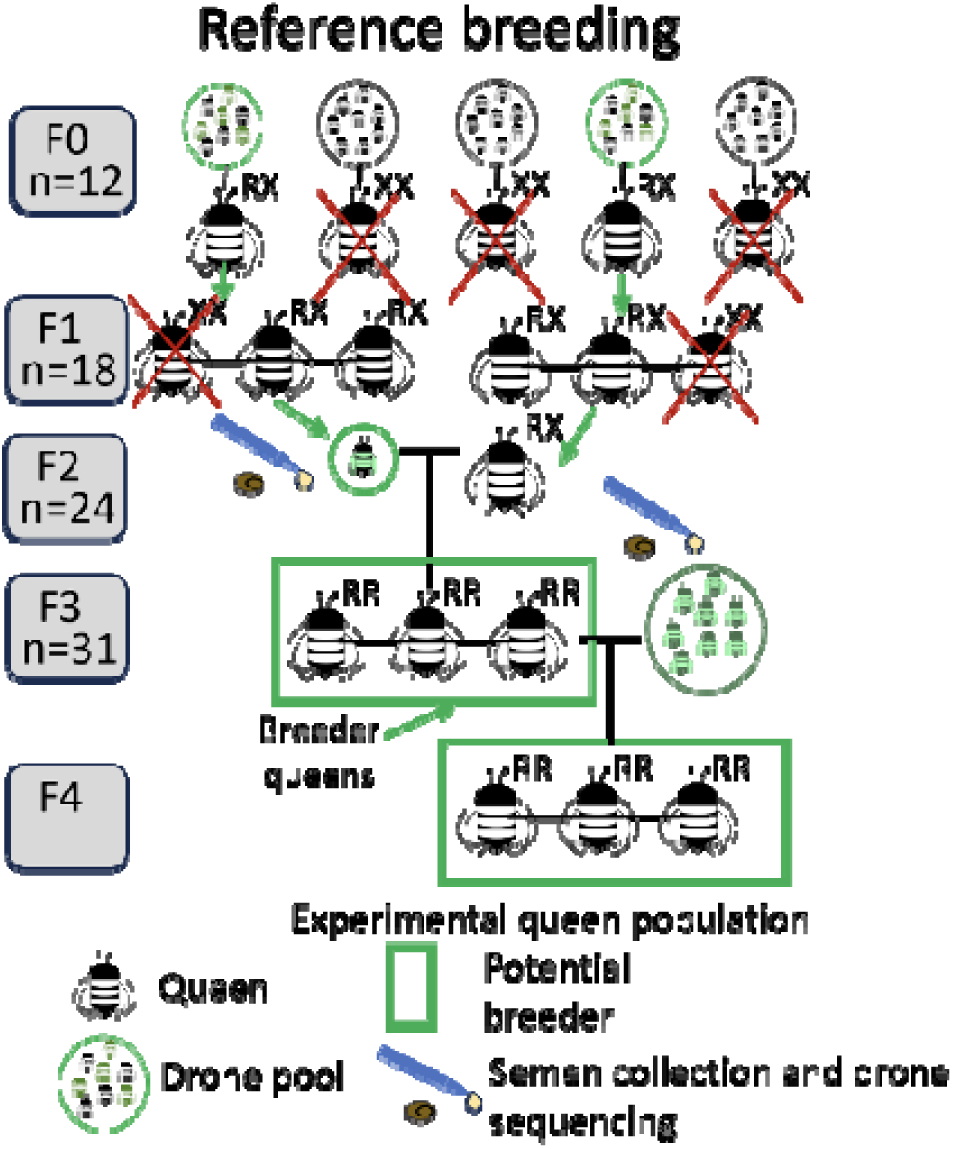
Breeding scheme for reference stocks. F0 and F1 queen genotypes are identified by sequencing their haploid drone sones. F1 and F2 daughters were reared from queen with a reference allele. F2 queens were genotyped via Sanger sequencing and inseminated with semen for a single drone. This was repeated for F3. The experimental colonies will be produced by inseminating the queens homozygous for the reference allele with pooled semen from drones produced by a homozygous mother. N for each generation signifies the number of queens with the desired genotype for each generation.

## Discussion

Our results represent the first successful selection of honey bees using MAS based on AI-predicted protein structure. We have established a breeding population of honey bees homozygous for the reference Vg variant. This proof of concept demonstrates the potential to employ any Vg variant in our scheme. We are therefore continuing our success by breeding for a Vg variant with predicted structural differences likely to impact the protein’s lipid-binding capacity. The functional effects of possessing this variant will be subsequently compared to that of the reference variant both biochemically, physiologically, and at the colony level in the field.

MAS in most other economically important species has resulted in significant economic benefits both to breeders and farmers and ranchers (Gutiérrez-Reinoso et al., 2023; Moreau et al., 2000; Peace, 2017; Strabel, 2024; Yu et al., 2000). The economic drawbacks from the lack of progress in developing successful MAS are felt acutely in the honey bee industry, especially as the biology of honey bees increases the difficulty and decreases the efficiency of trait-based selection. First, complications arise because the collective phenotypes of honey bee colonies emerge from the social interplay of traits of worker bees, which are functionally sterile females. As such, the individuals most directly responsible for the valuable colony traits can never actually be bred. Thus, current honey bee breeding programs rely on the screening of many (often hundreds of) colonies to assess and score the social traits of interest. The best-scoring colonies are used for production of virgin queens and male drones for controlled free-flight mating or instrumental insemination, and successfully mated queens are assisted in starting new colonies. The traits of these colonies can be scored after several months or more when the queens’ offspring make up the colony population, and the process is repeated. The trait screening most widely adopted by commercial queen breeders in the United States is colony hygienic testing developed at the University of Minnesota (Spivak, 1996). Hygienic behavior is the uncapping and removal of diseased or parasitized brood. To test for this trait, a breeder would freeze-kill a section of brood using liquid nitrogen. Twenty-four hours later, the trait is scored by determining the percentage of freeze-killed brood that has been removed from the colony – i.e. as a group effort. While the protocol is relatively straightforward, it is also time consuming, involves handling of hazardous liquid nitrogen, and requires financial investment into labor. Other selection programs, especially in for Varroa-resistance traits, are even more onerous, putting screenings out of reach for most of the industry.

Successful development of MAS for honey bees will have economic benefits across the industry. For breeders, MAS reduces generation time as selection can occur at very young ages. For most desirable traits in honey bees, phenotypic selection relies on rearing many queens and screening their resulting, mature colonies after a full field season to identify potential breeder queens. MAS allows these potential breeding individuals to be screened as early as adult emergence, reducing generation time from up to a full year to just under a month. This early screening also allows breeders to drastically reduce the number of colonies that they must manage, even as the number of potential breeders to be evaluated can increase, as rejected breeder queens can be culled before they are introduced into a colony. As such, breeders need only to maintain enough stock to sustain their breeding operation and avoid inbreeding. This further reduces costs for equipment, labor, and colony management.

For beekeepers, successful and rapid MAS will ideally allow them to reduce colony losses overtime as breeders are able to address new challenges as they arise. As most commercial beekeepers requeen their colonies every one to two years, introducing new lines of selected queens would fit seamlessly into their established practices. The reduction of colony losses would also reduce the need to aggressively “split” colonies in the spring to make up their numbers. Colonies with strong populations are better able to thrive through dearth periods and resist disease (Harbo, 1986; Spivak and Gilliam, 1993), which could help to reduce the rising colony losses occurring during the summers (Agostina Giacobino et al., 2024). The introduction of feed-efficient and disease resistant queens would also reduce costs in terms of supplemental feeding, treatments, and the labor required to perform management.

Honey bee researchers have been working to develop breeding markers to address these difficulties. Prior efforts have identified DNA markers for hygienic behavior (Lapidge et al., 2002), Varroa sensitive hygiene (Tsuruda et al., 2012), and foraging behavior (Rüppell et al., 2004). These markers, however, show that each trait is influenced by multiple, often interacting genome regions, each exerting only moderate effects. Because of this complexity, there has been very little functional testing of genes or gene products from candidate genome regions, and almost none in an applied, apicultural context (Sainsbury et al., 2022; Wang et al., 2010). The most successful testing to date is an intensive effort using proteomic molecular profiles to identify peptide markers for hygienic behavior (Guarna et al., 2017). MAS on the peptides was effective in increasing colony hygienic behavior. However, this exciting achievement relies on proteomic screening technology that is fast-evolving, highly technological, expensive, and expertise- and instrument-driven. These factors result in high barriers to adoption by the bee breeding industry.

In this paper, we present a single-gene-target MAS approach that is fast, cost-effective, and overcomes financial and time-consuming challenges without relying on complex technical methods, hazardous materials, or labor-intensive procedures. For our approach most costs rest in the initial long-read sequencing of whole genes to identify predicted high value variants. At present, this research and development would likely still need to be performed by scientists familiar with gene function and protein structure. For those already established, simple Sanger sequencing can be used to screen potential breeder queens. Costs for Sanger sequencing are low, typically less than $5-10 per sample. The outputs are also relatively simple to interpret, with sequence alignments to desired, established target variants requiring only cutting and pasting into freely available online tools. These aspects make this approach accessible to queen breeders with minimal training and investment relative to the costs associated with traditional trait-based colony screening.

While this approach can easily be expanded to include several targets, the cost-effectiveness relies on the ability to detect signatures for the desired protein variants. This may require using several signature elements or exclude colonies that contains individuals with rare SNPs. For example, design of the signature target for the lipid-variant identified two drones with a rare SNPs obstructing the signature signal. By simply excluding the colony from our starting material we were able to perfectly separate the target variant from the remaining haplotypes using the signature. Further, with the high rate of recombination in the honey bee genome, this approach relies that little to no recombination occurs in the targeted variants which might disrupt the signatures. Because of this, periodic long-range sequencing checks in the breeding populations may be required.

The success of our initiative will provide a cost-effective and readily adaptable approach to MAS in honey bees. Our proof of concept breeding for Vg protein variants is rooted in the substantial literature supporting a role for Vg in many health promoting pathways and leverages the extensive natural genetic diversity recently discovery for honey bee Vg. Our approach can lead directly to the use of MAS to target nutritional and health beneficial Vg variants, for example those predicted enhanced lipid- and zinc-binding efficiency-traits desirable for both queen breeders and beekeepers. Also, the effective screening and the use of cutting-edge AI-driven algorithms to predict the effects of genetic variants enables a quick turn-around if and when new threats arise from pests and pathogens. Our concept represents a new and promising approach to solving the challenge of breeding honey bees to provide sustainable pollination services for the future.

## Acknowledgements

Nanopore sequencing was performed by the Centre for Integrative Genetics (CIGENE), Norwegian University of Life Sciences and Sanger sequencing was performed by the DNA Services Labs, University of Illinois at Urbana-Champaign. The authors acknowledge The Research Council of Norway (RCN) grant number 335244 for funding toward running costs and positions and RCN grant number 350231 for travel grant and conference support. Mention of trade names or commercial products in this publication is solely for the purpose of providing specific information and does not imply recommendation or endorsement by the U.S. Department of Agriculture. U.S. Department of Agriculture is an equal opportunity provider and employer.

## Author Contribution Statements

Conceptualization GVA, KI; Data curation VL, KI; Formal analysis VL, GVA, SO, EP, GD, KI; Funding acquisition GVA, KI; Investigation VL, GVA, SO, EP, GD, KI; Methodology VL, GVA, SO, EP, GD, KI; Project administration GVA, KI; Resources GVA, KI; Software VL, KI; Supervision VL, GVA, KI; Validation; Visualization VL, KI; Writing—original draft VL, KI; Writing—review & editing VL, GVA, SO, EP, GD, KI.

Authors have no conflict of interest.

