## Supplementary Material for "AI-Enhanced Marker-Assisted Selection Concept for The Multifunctional Honey Bee (Hymenoptera: Apidea) Protein Vitellogenin (Vg)"

```
                             369
Vg_Exon2(ID:_406088)         cggggacagtttcagccgacttccagcacaattggcaagtcgggaatgagtacacgtatctcgttcgaagtcgcacgttgacgagtctgggcgatctgtcggacgtgcataccggtattctgataaaggcgttgctcaccgtacag
Exon_2_hap_1_(20_drones)     cggggacagtttcagccgacttccagcacaattggcaagtcgggaatgagtacacgtatctcgttcgaagtcgcacgttgacgagtctgggcgatctgtcggacgtgcataccggtattctgataaaggcgttgctcaccgtacag
Exon_2_hap_2_(17_drones)     cggggacagtttcagccgacttccagcacaattggcaagtcgggaatgagtacacgtatctcgttcgaagtcgcacgttgacgagtctgggcgatctgtcggacgtgcataccggtattctgataaaggcgttgctcaccgtacag
Exon_2_hap_3_(28_drones)     cggggacagtttcagccgacttccagcacaattggcaagtcgggaatgagtacacgtatctcgttcgaagtcgcacgttgacgagtctgggcgatctgtcggacgtgcataccggtattctgataaaggcgttgctcaccgtacag
Exon_2_hap_4_(13_drones)     cggggacagtttcagccgacttccagcacaattggcaagtcgggaatgagtacacgtatctcgttcgaagtcgcacgttgacgagtctgggcgatctgtcggacgtgcataccggtattctgataaaggcgttgctcaccgtacag
Exon_2_hap_5_(3_drones)      cggggacagtttcagccgacttccagcacaattggcaagtcgggaatgagtacacgtatctcgttcgaagtcgcacgttgacgagtctgggcgatctgtcggacgtgcataccggtattctgataaaggcgttgctcaccgtacag
Exon_2_hap_6_(30_drones)     cggggacagtttcagccgacttccagcacaattggcaagtcgggaatgagtacacgtatctcgttcgaagtcgcacgttgacgagtctgggcgatctgtcggacgtgcataccggtattctgataaaggcgttgctcaccgtacag
Exon_2_hap_7_(30_drones)     cggggacagtttcagccgacttccagcacaattggcaagtcgggaatgagtacacgtatctcgttcgaagtcgcacgttgacgagtctgggcgatctgtcggacgtgcataccggtattctgataaaggcgttgctcaccgtacag

                                        526                                             574
Vg_Exon2(ID:_406088)         gcgaaggattcaaacgtgcttgccgccaaagtgtggaacggccaatatgctcgtgtccaacagtccatgcctgacggatgggaaacggagatctccgaccaaatgttggagctccgtgacctgcctatttccggcaaaccgttcca
Exon_2_hap_1_(20_drones)     gcgaaggattcaaacgtgcttgccgccaaagtgtggaacggccaatatgctcgtgtccaacagtccatgcctgacggatgggaaacggagatctccgaccaaatgttggagctccgtgacctgcctatttccggcaaaccgttcca
Exon_2_hap_2_(17_drones)     gcgaaggattcgaacgtgcttgccgccaaagtgtggaacggccaatatgctcgtgtccagcagtccatgcctgacggatgggaaacggagatctccgaccaaatgttggagctccgtgacctgcctatttccggcaaaccgttcca
Exon_2_hap_3_(28_drones)     gcgaaggattcgaacgtgcttgccgccaaagtgtggaacggccaatatgctcgtgtccagcagtccatgcctgacggatgggaaacggagatctccgaccaaatgttggagctccgtgacctgcctatttccggcaaaccgttcca
Exon_2_hap_4_(13_drones)     gcgaaggattcgaacgtgcttgccgccaaagtgtggaacggccaatatgctcgtgtccagcagtccatgcctgacggatgggaaacggagatctccgaccaaatgttggagctccgtgacctgcctatttccggcaaaccgttcca
Exon_2_hap_5_(3_drones)      gcgaaggattcgaacgtgcttgccgccaaagtgtggaacggccaatatgctcgtgtccagcagtccatgcctgacggatgggaaacggagatctccgaccaaatgttggagctccgtgacctgcctatttccggcaaaccgttcca
Exon_2_hap_6_(30_drones)     gcgaaggattcaaacgtgcttgccgccaaagtgtggaacggccaatatgctcgtgtccaacagtccatgcctgacggatgggaaacggagatctccgaccaaatgttggagctccgtgacctgcctatttccggcaaaccgttcca
Exon_2_hap_7_(30_drones)     gcgaaggattcaaacgtgcttgccgccaaagtgtggaacggccaatatgctcgtgtccaacagtccatgcctgacggatgggaaacggagatctccgaccaaatgttggagctccgtgacctgcctatttccggcaaaccgttcca

                                                                    700          714                                                       773
Vg_Exon2(ID:_406088)         gatcaggatgaaacacggtttgatcagggatctgatcgttgatcgcgacgtccccacttgggaggtgaacatactcaagagcatcgttggccaactgcaagtggacactcaaagtgaaaacgccgtgaaggtgaacagcgtgcaag
Exon_2_hap_1_(20_drones)     gatcaggatgaaacacggtttgatcagggatctgatcgttgatcgcgacgtccccacttgggaggtgaacatactcaagagcatcgttggccaactgcaagtggacactcaaagtgaaaacgccgtgaaggtgaacagcgtgcaag
Exon_2_hap_2_(17_drones)     gatcaggatgaaacacggtttgatcagggatctgatcgtggatcgcgacgtccccacttgggaggtgaacatactcaagagcatcgttggccaactgcaagtggacactcaaggtgaaaacgccgtgaaggtgaacagcgtgcaag
Exon_2_hap_3_(28_drones)     gatcaggatgaaacacggtttgatcagggatctgatcgttgatcgcgacgtcctcacttgggaggtgaacatactcaagagcatcgttggccaactgcaagtggacactcaaggtgaaaacgccgtgaaggtgaacagcgtgcaag
Exon_2_hap_4_(13_drones)     gatcaggatgaaacacggtttgatcagggatctgatcgttgatcgcgacgtccccacttgggaggtgaacatactcaagagcatcgttggccaactgcaagtggacactcaaggtgaaaacgccgtgaaggtgaacagcgtgcaag
Exon_2_hap_5_(3_drones)      gatcaggatgaaacacggtttgatcagggatctgatcgttgatcgcgacgtccccacttgggaggtgaacatactcaagagcatcgttggccaactgcaagtggacactcaaggtgaaaacgccgtgaaggtgaacagcgtgcaag
Exon_2_hap_6_(30_drones)     gatcaggatgaaacacggtttgatcagggatctgatcgttgatcgcgacgtccccacttgggaggtgaacatactcaagagcatcgttggccaactgcaagtggacactcaaggtgaaaacgccgtgaaggtgaacagcgtgcaag
Exon_2_hap_7_(30_drones)     gatcaggatgaaacacggtttgatcagggatctgatcgttgatcgcgacgtccccacttgggaggtgaacatactcaagagcatcgttggccaactgcaagtggacactcaaggtgaaaacgccgtgaaggtgaacagcgtgcaag

                                                                                                                                                     946
Vg_Exon2(ID:_406088)         ttccgaccgacgacgagccgtacgcctcgttcaaagccatggaggactccgtgggagggaaatgcgaggttctctacgatatagcgcctttgtcggactttgtgatccacagatcgccggaattggtgccgatgccgacgttgaag
Exon_2_hap_1_(20_drones)     ttccgaccgacgacgagccgtacgcctcgttcaaagccatggaggactccgtgggagggaaatgcgaggttctctacgatatagcgcctttgtcggactttgtgatccacagatcgccggaattggtgccgatgccgacgttgaag
Exon_2_hap_2_(17_drones)     ttccgaccgacgacgagccgtacgcctcgttcaaagccatggaggactccgtgggagggaaatgcgaggttctctacgatatagcgcctttgtcggactttgtgatccacagatcgccggaattggtgccgatgccgacgttgaag
Exon_2_hap_3_(28_drones)     ttccgaccgacgacgagccgtacgcctcgttcaaagccatggaggactccgtgggagggaaatgcgaggttctctacgatatagcgcctttgtcggactttgtgatccacagatcgccggaattggtgccgatgccgacgttgaag
Exon_2_hap_4_(13_drones)     ttccgaccgacgacgagccgtacgcctcgttcaaagccatggaggactccgtgggagggaaatgcgaggttctctacgatatagcgcctttgtcggactttgtgatccacagatcgccggaattggtgccgatgccgacttgaag
Exon_2_hap_5_(3_drones)      ttccgaccgacgacgagccgtacgcctcgttcaaagccatggaggactccgtgggagggaaatgcgaggttctctacgatatagcgcctttgtcggactttgtgatccacagatcgccggaattggtgccgatgccgacgttgaag
Exon_2_hap_6_(30_drones)     ttccgaccgacgacgagccgtacgcctcgttcaaagccatggaggactccgtgggagggaaatgcgaggttctctacgatatagcgcctttgtcggactttgtgatccacagatcgccggaattggtgccgatgccgacgttgaag
Exon_2_hap_7_(30_drones)     ttccgaccgacgacgagccgtacgcctcgttcaaagccatggaggactccgtgggagggaaatgcgaggttctctacgatatagcgcctttgtcggactttgtgatccacagatcgccggaattggtgccgatgccgacgttgaag

                                        964                              997                                      1039                                  1087
Vg_Exon2(ID:_406088)         ggcgatggccgccacatggaggtgatcaagatcaagaacttcgataactgcgatcaaaggataaattatcatttcggtatgacggataactcgaggttggaacctggaacgaacaagaatggaaagttcttctcg
Exon_2_hap_1_(20_drones)     ggcgatggccgccacatggaggtgatcaagatcaagaacttcgataactgcgatcaaaggataaattatcatttcggtatgacggataactcgaggttggaacctggaacgaacaagaatggaaagttcttctcg
Exon_2_hap_2_(17_drones)     ggcgatggccgccacatggaggtgatcaagatcaagaacttcgacaactgcgatcaaaggataaattatcatttcggtatgacggacaactcgaggttggaacctggaacgaacaagaatggaaagttcttctcg
Exon_2_hap_3_(28_drones)     ggcgatggccgtcacatggaggtgatcaagatcaagaacttcgacaactgcgatcaaaggataaattatcatttcggtatgacggacaactcgaggttggaacctggaacgaacaagaatggaaagttcttctcg
Exon_2_hap_4_(13_drones)     ggcgatggccgccacatggaggtgatcaagatcaagaacttcgataactgcgatcaaaggataaattatcatttcggtatgacggataactcgaggttggaacctggaacgaacaagaatggaaagttcttctcg
Exon_2_hap_5_(3_drones)      ggcgatggccgccacatggaggtgatcaagatcaagaacttcgataactgcgatcaaaggataaattatcatttcggtatgacggataactcgaggttggaacctggaacgaacaagaatggaaagttcttctcg
Exon_2_hap_6_(30_drones)     ggcgatggccgccacatggaggtgatcaagatcaagaacttcgataactgcgatcaaaggataaattatcatttcggtatgacggataactcgaggttggaacctggaacgaacaagaatggaaagttcttctcg
Exon_2_hap_7_(30_drones)     ggcgatggccgtcacatggaggtgatcaagatcaagaacttcgacaactgcgatcaaaggataaattatcatttcggtatgacggacaactcgaggttggaacctggaacgaacaagaatggaaagttcttctcg
```

*Figure S1: Exon 2 of Vg (line 1, nucleotide 369-1087) aligned to the unique haplotypes of exon 2 identified in the Pol-line drones (hap 1-7 on line 2-8). The 9 identified SNPs are in bold, and the reference nucleotide is marked in green, while the alternative nucleotide is marked in yellow.*

```
                              2085                                                  2894                                      2927                2945
Vg_Exon_4_(ID:406088)         gtgaaccaccttctcacgcatcacgaatacgactacgaactctcaaggggatacatcgacgagaagattttggagaaccagaacatcatcacccatatgatcctcaattacgtgggaagcgaagacagcgtgatcccgcgcatcct
Exon_4_hap_1_(74_drones)      gtgaaccaccttctcacgcatcacgaatacgactacgaactctcaaggggatacatcgacgagaagattttggagaaccagaacatcatcacccatatgatcctcaattacgtgggaagcgaagacagcgtgatcccgcgcatcct
Exon_4_hap_2_(17_drones)      gtgaaccaccttctcacgcatcacgaatacgactacgaactctcaaggggatacatcgacgagaagattttggagaaccagaacatcatcacccacatgatcctcaattacgtgggaagcgaagacagcgtgatcccgcgcatcct
Exon_4_hap_3_(48_drones)      gtgaaccaccttctcacgcatcacgaatacgactacgaactctctcggggatacatcgacgagaagattttggagaatcagaacatcatcacccacatgatcctcaattacgtgggaagcgaagacagcgtgatcccgcgcatcct
Exon_4_hap_3_(2_drones)       gtgaaccaccttctcacgcatcacgaatacgactacgaactctctcggggatacatcgacgagaagattttggagaatcagaacatcatcacccacatgatcctcaattacgtgggaagcgaagacagcgtgatcccgcgcatcct

Vg_Exon_4_(ID:406088)         ctaccttacctggtactcctccaacggcgacataaaagtaccttccaccaaagtgctagccatgatctcgagcgtgaaatcattcatggagttgagcctgaggagcgtgaaggaccgagaaacgattatttcggcggccgagaaga
Exon_4_hap_1_(74_drones)      ctaccttacctggtactcctccaacggcgacataaaagtaccttccaccaaagtgctagccatgatctcgagcgtgaaatcattcatggagttgagcctgaggagcgtgaaggaccgagaaacgattatttcggcggccgagaaga
Exon_4_hap_2_(17_drones)      ctaccttacctggtactcctccaacggcgacataaaagtaccttccaccaaagtgctagccatgatctcgagcgtgaaatcattcatggagttgagcctgaggagcgtgaaggaccgagaaacgattatttcggcggccgagaaga
Exon_4_hap_3_(48_drones)      ctaccttacctggtactcctccaacggcgacataaaagtaccttccaccaaagtgctagccatgatctcgagcgtgaaatcattcatggagttgagcctgaggagcgtgaaggaccgagaaacgattatttcggcggccgagaaga
Exon_4_hap_3_(2_drones)       ctaccttacctggtactcctccaacggcgacataaaagtaccttccaccaaagtgctagccatgatctcgagcgtgaaatcattcatggagttgagcctgaggagcgtgaaggaccgagaaacgattatttcggcggccgagaaga

                                                                                                       3209     3218                     3242                   3266
Vg_Exon_4_(ID:406088)         tcgccgaggagttgaagatcgtccccgaagagctcgttcctctggaaggaaacttgatgataaacaacaaatatgcgttgaaattcttccccttcgataaacacattctcgacaaattacccacg
Exon_4_hap_1_(74_drones)      tcgccgaggagttgaagatcgtccccgaagagctcgttcctctggaaggaaacttgatgataaacaacaaatatgcgttgaaattcttccccttcgataaacacattctcgacaaattacccacg
Exon_4_hap_2_(17_drones)      tcgccgaggagttgaagatcgtccccgaagagctcgttcctctggaaggaaacttgatgataaacaacaaatatgcgttgaaattcttccccttcgataagcacattctcgacaaattacccacg
Exon_4_hap_3_(48_drones)      tcgccgaggagttgaagatcgtccccgaagagctcgttcctctggaaggaaacttgatgataaacaataaatatgccttgaaattcttccccttcgataagcacattctcgacaaattacccacg
Exon_4_hap_3_(2_drones)       tcgccgaggagttgaagatcgtccccgaagagctcgttcctctggaaggaaacttgatgataaacaataaatatgccttgaaattcttccccttcgataagcacattctcgacaaattacccacg
```

*Figure S2: The middle and end part of exon 4 of Vg (line 1, nucleotide 2085-3266) aligned to the unique haplotypes of the same section in exon 4 identified in the Pol-line drones (hap 1-3 on line 2-5). The 6 identified SNPs are in bold, and the reference nucleotide is marked in green, while the alternative nucleotide is marked in yellow. Haplotype 3 was found for 50 drones in total; 48 represent Vg variant 15, while 2 represent Vg variant 7 (line 4 and 5, respectively).*

**Table S1** **The 26 Vg variants**

| Vg variants | Number of drones | nsSNPs | | | | | | | | | | | | | |
|---|---|---|---|---|---|---|---|---|---|---|---|---|---|---|---|
| 1 | 24 | no nsSNPs | | | | | | | | | | | | | |
| 2 | 22 | p.S146G | | | | | | | | | | | | | |
| 3 | 1 | p.S146G | p.S154R | | | | | | | | | | | | |
| 4 | 1 | p.S146G | p.S154E | | | | | | | | | | | | |
| 5 | 7 | p.S146G | p.S1110T | | | | | | | | | | | | |
| 6 | 4 | p.S146G | p.V1451A | p.T1503A | | | | | | | | | | | |
| 7 | 2 | p.S146G | p.S1110T | p.N1220S | p.R1284K | | | | | | | | | | |
| 8 | 5 | p.S146G | p.S1110T | p.L1291I | p.R1292S | | | | | | | | | | |
| 9 | 1 | p.S146G | p.S1110T | p.R1292S | p.V1451A | | | | | | | | | | |
| 10 | 45 | p.T6M | p.P126L | p.S146G | p.S1110T | p.T1503A | | | | | | | | | |
| 11 | 2 | p.S146G | p.S1110T | p.L1291I | p.R1292S | p.V1451A | | | | | | | | | |
| 12 | 15 | p.S146G | p.S1110T | p.L1291I | p.R1292S | p.V1451A | p.T1503A | | | | | | | | |
| 13 | 4 | p.S146G | p.S1110T | p.N1220S | p.R1284K | p.I1398V | p.V1451A | p.T1503A | | | | | | | |
| 14 | 21 | p.S146G | p.S1110T | p.R1284K | p.I1398V | p.V1451A | p.T1503A | p.I1536V | | | | | | | |
| 15 | 92 | p.S146G | p.S1110T | p.N1220S | p.R1284K | p.I1398V | p.V1451A | p.T1503A | p.I1536V | | | | | | |
| 16 | 6 | p.S146G | p.S1110T | p.N1220S | p.L1291I | p.R1292S | p.V1451A | p.T1503A | p.I1536V | | | | | | |
| 17 | 21 | p.S1110T | p.V1199I | p.N1220S | p.R1284K | p.I1398V | p.V1451A | p.T1503A | p.I1536V | | | | | | |
| 18 | 5 | p.S15A | p.G25E | p.S146G | p.N326S | p.S1110T | p.N1220S | p.I1398V | p.T1503A | p.L1751F | | | | | |
| 19 | 17 | p.S146G | p.S375L | p.T522I | p.V538I | p.S1110T | p.N1220S | p.I1398V | p.V1451A | p.T1503A | | | | | |
| 20 | 2 | p.S146G | p.S803N | p.S1110T | p.N1220S | p.R1284K | p.I1398V | p.V1451A | p.T1503A | p.I1536V | | | | | |
| 21 | 2 | p.S146G | p.S1110T | p.R1174K | p.N1220S | p.R1284K | p.I1398V | p.V1451A | p.T1503A | p.I1536V | | | | | |
| 22 | 11 | p.S146G | p.S1110T | p.N1220S | p.R1284K | p.I1398V | p.V1451A | p.T1503A | p.I1536V | p.T1676S | | | | | |
| 23 | 4 | p.S146G | p.S1110T | p.M1159I | p.N1220S | p.L1291I | p.R1292S | p.I1398V | p.V1451A | p.T1503A | p.I1536V | | | | |
| 24 | 1 | p.S146G | p.S1110T | p.R1174K | p.N1220S | p.R1284K | p.V1451A | p.T1503A | p.I1536V | p.T1676S | p.G1678S | | | | |
| 25 | 11 | p.S146G | p.S1110T | p.N1220S | p.R1284K | p.I1398V | p.V1451A | p.T1503A | p.I1536V | p.T1676S | p.G1678S | | | | |
| 26 | 4 | p.S15A | p.S146G | p.T311M | p.D608E | p.V952I | p.S1110T | p.N1220S | p.V1311M | p.T1503A | p.V1510L | p.I1536V | p.V1610M | p.L1670S | p.T1716M |
| Total: | **330** | | | | | | | | | | | | | | |
